# The role of synergy and antagonism in designing multidrug adaptive chemotherapy schedules

**DOI:** 10.1101/2020.05.13.094375

**Authors:** P.K. Newton, Y. Ma

## Abstract

Chemotherapeutic resistance via the mechanism of competitive release of resistant tumor cell subpopulations is a major problem associated with cancer treatments and one of the main causes of tumor recurrence. Often, chemoresistance is mitigated by using multidrug schedules (two or more combination therapies) that can act synergistically, additively, or antagonistically on the heterogeneous population of cells as they evolve. In this paper, we develop a three-component evolutionary game theory model to design two-drug adaptive schedules (timing and dose levels associated with *C*_1_(*t*) and *C*_2_(*t*)) that mitigate chemoresistance and delay tumor recurrence in an evolving collection of tumor cells with two resistant subpopulations: *R*_1_ (sensitive to drug 1, resistant to drug 2), and *R*_2_ (sensitive to drug 2, resistant to drug 1). A key parameter, *e*, takes us from synergistic (*e >* 0), to additive (*e* = 0), to antagonistic (*e <* 0) drug interactions. In addition to the two resistant populations, the model includes a population of chemosensitive cells, *S* that have higher baseline fitness but are not resistant to either drug. Using the nonlinear replicator dynamical system with a payoff matrix of Prisoner’s Dilemma (PD) type (enforcing a cost to resistance), we investigate the nonlinear dynamics of the three-component system (*S, R*_1_*, R*_2_), along with an additional tumor growth model whose growth rate is a function of the fitness landscape of the tumor cell populations. We show that *antagonistic* drug interactions generally result in slower rates of adaptation of the resistant cells than *synergistic* ones, making them more effective in combating the evolution of resistance. We then design closed loops in the three-component phase space by shaping the fitness landscape of the cell populations (i.e. altering the evolutionary stable states of the game) using appropriately designed time-dependent schedules (adaptive therapy), altering the dosages and timing of the two drugs using information gleaned from constant dosing schedules. We show that the bifurcations associated with the evolutionary stable states are transcritical, and we detail a typical antagonistic bifurcation that takes place between the sensitive cell population *S* and the *R*_1_ population, and a synergistic bifurcation that takes place between the sensitive cell population *S* and the *R*_2_ population for fixed values of *C*_1_ and *C*_2_. These bifurcations help us further understand why antagonistic interactions are more effective at controlling competitive release of the resistant population than synergistic interactions in the context of an evolving tumor.

## I. INTRODUCTION

We study a mathematical model to explore the role of synergisitic vs. antagonistic multidrug interactions on an evolving population of cancer cells in a tumor. Our model builds on the (single drug) adaptive therapy model developed in [1] and is based on a replicator dynamical system of three (well-mixed) populations of cells: (i) sensitive cancer cells, *S*, that are sensitive to both drug 1 and drug 2, (ii) a resistant population, *R*_1_, that is sensitive to drug 1 but resistant to drug 2, and (iii) resistant population, *R*_2_, that is sensitive to drug 2 but resistant to drug 1. The replicator dynamical system governing the relative frequencies of (*S, R*_1_*, R*_2_) makes use of a 3×3 payoff matrix *A* of Prisoner’s Dilemma type [2–5]. We introduce chemotherapeutic schedules using time-dependent dosing functions *C*_1_(*t*), *C*_2_(*t*) that alter the fitness levels of the sensitive cell population and the two resistant populations independently allowing us to shape the fitness landscape of the population of cells adaptively as the tumor evolves [6–13]. Our general goal in this context is to design schedules that delay tumor recurrence (re-growth) due to competitive release of the resistant cell population [14–17] and to quantify the role of synergistic and antagonistic drug interactions in this process.

To quantify the synergistic vs. antagonistic effects of the two toxins on the evolving populations, we use a parameter *e* which we alter. When *e >* 0, the drugs interact synergistically, when *e* = 0, the drugs are additive, and when *e <* 0, they interact antagonistically. Level curves of the fitness profiles as a function of *C*_1_ and *C*_2_ are shown in figure 1 which can be compared with figure 1 of [18]. Details of how we define our fitness functions in the context of our replicator model will be described in §II. In the neutral case, additive drug interactions (figure 1(b)) accomplish exactly the kill rate that the sum of each of the two would accomplish acting independently. By contrast, synergistic drug interactions (figure 1(a)) bend the curves inward, indicating that the growth rates (fitness) are lowered more than the two dosages would accomplish independently, while antagonistic interactions (figure 1(c)) bend the curves outward, indicating that the growth rates are lowered less than the two dosages would accomplish independently.

**FIG. 1.**
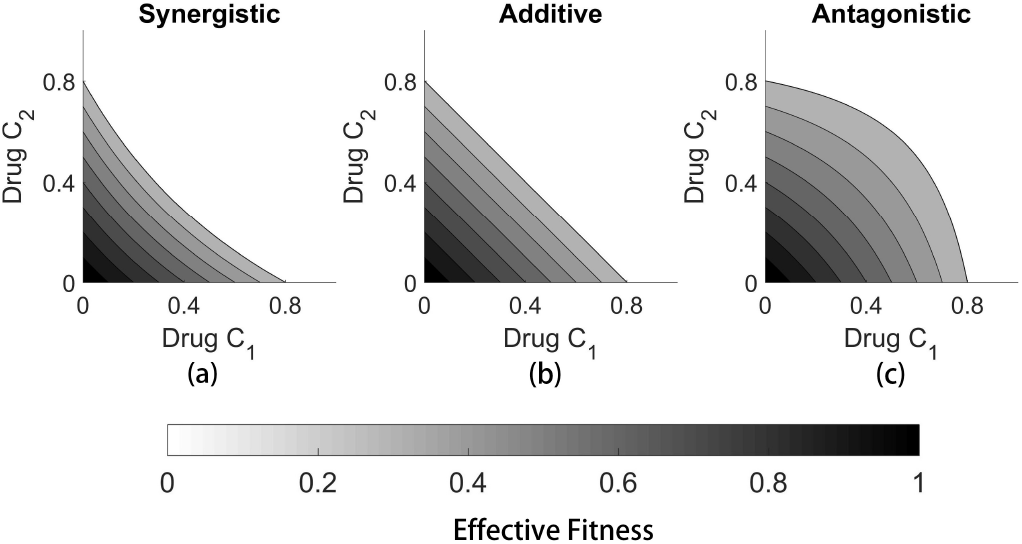
(a) *e >* 0 showing synergistic profiles; (b) *e* = 0 showing additive profiles; (c) *e <* 0 showing antagonistic profiles.

There is a large and dedicated literature on characterizing the interactions of many different combinations of toxins on static cell populations. As far as we are aware, the first comprehensive study of synergistic vs antagonistic effects was carried out by Bliss [19] in 1939, using joint probabilities, leading to a formula that is commonly called the Bliss index for drug interactions. A similar but slightly modified criterion was introduced by Loewe [20] and more recently developed further by Chou and collaborators [21]. These indices have all been used to help quantify the many different types of interactions that can occur with two or more toxins applied jointly in a static population of cells. In this context, it is common to assume that synergistic interactions are desirable in most circumstances, as a lower total dosage accomplishes the same kill rate as a higher dose would accomplish if the drugs acted independently. These kinds of studies have been used effectively to choose appropriate drug cocktails to individual patients by testing wide ranges of combinations on tissue samples obtained from patient tumors [22].

When the interacting population of cells are evolving, however, the relevant criteria become more complex. This is due to the fact that the subpopulations of cells respond differently to the different toxins applied, and as they respond, an ever-changing (adaptive) combination of toxins might be required to accomplish a given goal. Instead of necessarily killing the maximum number of cancer cells with the least amount of toxin, it is often the case that the goal becomes avoiding chemo-resistance and delaying unwelcome tumor recurrence to the maximum extent possible. A strategy called *resistance management* [23] is often advocated and occasionally implemented [24]. These kinds of strategies have been advocated and implemented in chemotherapy settings [7, 10, 11, 14, 25–27], but perhaps have been most elegantly and thoroughly carried out in a bacterial setting (since experiments are more practical) by Kishony and collaborators [18, 28–32] who even discuss strategies that might reverse antibiotic resistance [31]. See also [33] for recent work discussing both microbial populations and cancer cells and [34, 35] for novel sequential therapy methods. In the context of evolving microbial populations [36], mutations occur frequently and it is important to consider not only pre-existing mutated subpopulations, but also mutations that occur as a result of the application of antibiotic agents. In the case of chemotherapeutic resistance in tumors, it is often assumed that resistant mutations occured before the application of treatment, hence it is common to separate the subpopulations into sensitive and resistant subpopulations as we do in our deterministic model which does not include further mutations during treatment. See [37] for further discussions of these and related issues, and [38, 39] for discussions of the general approach of using evolutionary game theory in biology.

In section §II we introduce the details of the three-component replicator dynamical system that we use. Section §III describes the effects of constant chemotherapy schedules on the evolving populations, using the range of values 0 ≤*C*_1_ ≤1, 0 ≤ *C*_2_ ≤ 1 (with a total dose upper threshold *C*_1_ + *C*_2_ ≤ 1), along with our parameter *e* over a range of positive (synergistic) to negative (antagonistic) values. We describe in detail the transcritical bifurcations that occur and the tumor growth in response to the chemotoxins. In §IV we introduce adaptive time-dependent schedules (*C*_1_(*t*)*, C*_2_(*t*)) along with the parameter *e* with the goal of delaying tumor recurrence and we discuss the rate of adaptation of the cell populations in this context. Finally in §V we discuss the relevance of our model to the design of adaptive-therapy clinical trials.

## II. A THREE-COMPONENT REPLICATOR SYSTEM

### A. Why the prisoner’s dilemma game?

To understand why the prisoner’s dilemma (PD) game is a useful paradigm for tumor growth, consider a two-component system, with a population of healthy cells *H*, and a population of sensitive cancer cells *S*, each competing for the best payoffs. The standard version of the prisoner’s dilemma payoff matrix is:

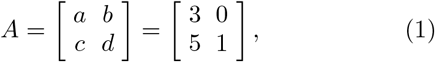

where the first row corresponds to my payoffs if I choose to cooperate (associated with the healthy cell population (H)) and the second row corresponds to my payoffs if I choose to defect (associated with chemo-sensitive cells (S)). I am better off defecting than cooperating if my opponent cooperates (first column), since my payoff is 5 instead of 3. I am also better off defecting than co-operating if my opponent defects (second column), since my payoff is 1 rather than 0. No matter what my opponent does, I am better off defecting. But my opponent uses the same logic and comes to the same conclusion - that she is also better off defecting, no matter what I do. The result is that we both defect and get a payoff of 1 (second row, second column) which is less than we would have gotten had we both cooperated (payoff of 3). The fact that defect-defect is a sub-optimal Nash equilibrium (neither player is better off by making a different choice) is the essence of the prisoner’s dilemma game [2, 3] and the reason it is so widely used as the basis for modeling the evolution of cooperation in many different contexts. For cancer cell modeling, the healthy cells are the cooperators and the cancer cells are the defectors. Unlike other contexts in which game theory is used, cells are not making strategic decisions, instead, their strategy is encoded in their reproductive prowess, and selection is frequency dependent. In any mixed population 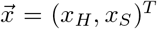, 0 ≤ *x_H_ ≤ 1; 0 ≤ x_S_ ≤ 1; x_H_ + x_S_* = 1, the fitness functions, 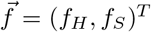, associated with the two subpopulations are:

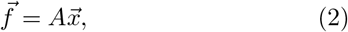

which in component form yields:

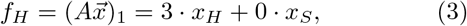

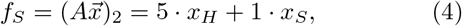

while the average fitness of the total population is given by the quadratic form:

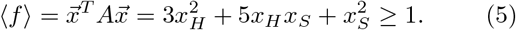

The average fitness of the healthy state (*x_H_, x_S_*) = (1, 0) is given by 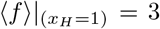, while that of the cancerous state (*x_H_, x_S_*) = (0, 1) is given by 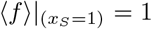, which minimizes the average fitness. For the static game, the cancerous state (*x_H_, x_S_*) = (0, 1) ≡ **p**^\**T*^ is a strict Nash equilibrium since **p**^\**T*^ *A* **p**^*^ *>* **p**^*T*^ A**p**^*^, for all **p**[2]. We can then embed this static game into an evolutionary context using the replicator dynamical system [3]:

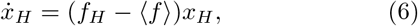

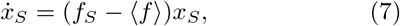

from which (using values from eqn (1)) it is straightfor-ward to show:

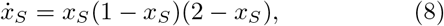

with fixed points at *x_S_* = 0, 1, 2. The cancerous state (*x_s_* = 1) then becomes an asymptotically stable fixed point of the dynamical system (8) and an evolutionary stable state (ESS) of the system (6), (7) which serves to drive the system to the strict Nash equilibrium under the flow. The fact that this ESS also corresponds to the one with the lowest average fitness is an extra feature of the PD game. For any initial condition containing at least one cancer cell: 0 *< x_S_* (0) ≤ 1, we have:

(i) *x_S_* → 1, *x_H_* → 0 as *t* → ∞
(ii) 〈*f*〉 → 1 as *t* → ∞.

The first condition (and the structure of the nonlinear equations) guarantees that the cancer cell population will saturate at the carrying capacity of 1 in an *S*-shaped (logistic) growth curve, while the second guarantees that this asmptotically stable carrying capacity is sub-optimal, since 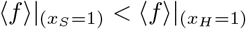. For these two reasons, the prisoner’s dilemma evolutionary game serves as a simple paradigm for tumor growth both in finite population models, as well as replicator system (infinite population) models [5, 25, 40, 41]. This two-component system alone, however, is not able to account for the evolution of resistance.

### B. The three-component model

The model we employ is a three-component replicator dynamical system for the three subpopulations of cells:(*S, R*_1_*, R*_2_) ≡ (*x*_1_*, x*_2_*, x*_3_)

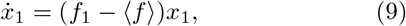

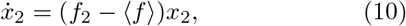

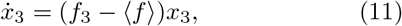

where each dependent variable represents the relative frequencies of cells, with *x*_1_+*x*_2_+*x*_3_ = 1. In these equations, *f_i_* represents the fitness of subpopulation *i* = 1, 2, 3, while *f* represents the average fitness of all three subpopulations. These equations then give rise to the obvious interpretation that if a given subpopulation’s fitness is above(below) the average, it grows(decays) exponentially - reproductive prowess is directly associated with the deviation of the fitness of a subpopulation from the average fitness of the entire population.

The fitness functions are frequency-dependent (i.e. non-constant), which couples eqns (9)-(11) nonlinearly. The subpopulation fitness *f_i_* (*i* = 1, 2, 3) function is given by:

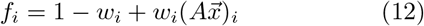

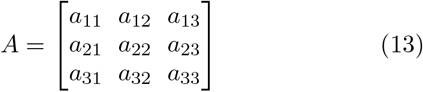

where *A* is a 3 × 3 payoff matrix which introduces the evolutionary game being played by *x*_1_, *x*_2_, and *x*_3_. In this paradigm, the sensitive population (*x*_1_) are the defectors (higher fitness) and both groups of resistant cells (*x*_2_ and *x*_3_) are the cooperators (lower, but not equal fitness). We use 0 ≤ *w_i_* ≤ 1 as a parameter to determine the strength of selection in the system. When *w_i_* ~ 0, selection is relatively weak and the evolutionary game does not play a big role in the balance of the three subpopulations. When *w_i_* ~ 1, selection is strong, and the game plays a bigger role. Both of those limiting cases have been discussed in the literature [42, 43], but the tumor response through the full range of values is poorly understood. We use the selection parameters *w_i_* as our mechanism to introduce chemotherapy, by way of altering the relative fitness of the three subpopulations:

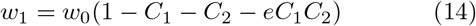

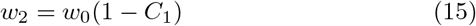

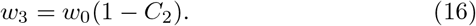

Here, *C*_1_ is the chemotherapy parameter associated with drug 1, *C*_2_ is that for drug 2, and *e* is our synergy (*e >* 0) vs. antagonism (*e <* 0) parameter, and we take *w*_0_ = 0.1 which sets the timescale in our simulations. Level curves of the effective fitness functions defined by eqns (14)-(16) are shown in figure 1. The average fitness of the population is given by:

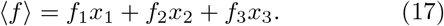

The condition for the payoff matrix *A* to be of (PD) type is:

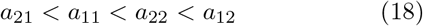

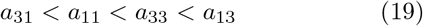

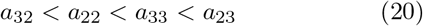

and for definiteness, we choose the specific values:

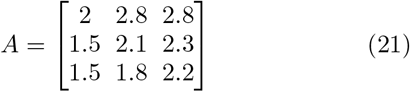

For the static game, it is easy to show that the cancerous state (*S, R*_1_*, R*_2_) = (*x*_1_*, x*_2_*, x*_3_) = (1, 0, 0) ≡ **p**^\**T*^ is a strict Nash equilibrium in the absence of chemotherapy (*C*_1_ = 0*, C*_2_ = 0) and an ESS for the replicator system, as shown in the diagram of figure 2(a) where the entire triangular region is the basin of attraction for the *S* population.

**FIG. 2.**
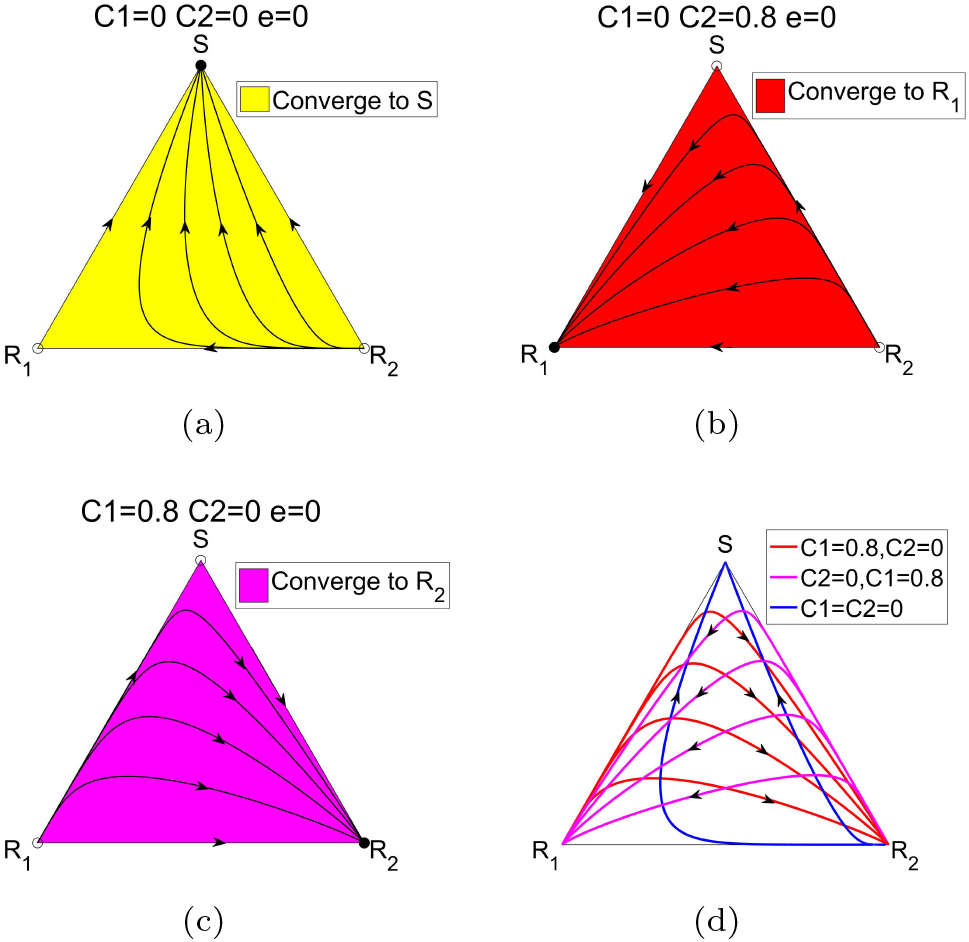
Depiction of evolutionary stable states (ESS) for different chemotherapy values. (a) With no chemotherapy, the tumor saturates to the *S* corner regardless of the initial make-up of the three subpopulations. (b) With *C*_1_ = 0, and *C*_2_ = 0.8, competitive release of the resistant population *R*_1_ drives all trajectories to the *R*_1_ corner. (c) With *C*_1_ = 0.8 and *C*_2_ = 0, competitive release of the resistant population *R*_2_ drives all trajectories to the *R*_2_ corner. (d) Trajectories associated with three different constant combinations of *C*_1_ and *C*_2_, depicting the overlap of the trajectories at different times.

A seperate important ingredient in our model is our tumor-growth equation, which we take as:

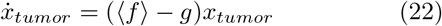

where the growth(decay) of the tumor is a function of the average fitness associated with the tumor minus a constant background fitness level *g*, associated with the surrounding tissue (say healthy cells) and micro-environment [44]. When the average fitness level of the population of cancer cells is higher than *g*, the tumor grows, and when it is lower, it regresses. Our chemotherapy functions *C*_1_ and *C*_2_ largely control this complex dynamic by modifying the Nash equilibria and ESS’s of the system via the fitness function (12).

## III. CONSTANT CHEMOTHERAPY

First, we examine the dynamical system for different constant levels of the chemo-parameters *C*_1_, *C*_2_ (0 ≤ *C*_1_ ≤ 1; 0 ≤ *C*_2_ ≤ 1; *C*_1_ + *C*_2_ ≤ 1) in the case of additive *e* = 0 interations, synergistic (*e >* 0) interactions, and antagonistic (*e <* 0) interactions.

### A. Additive interactions *e* = 0

Figure 2 shows the panel of trajectories for three different scenarios. To start, figure 2(a) shows the trajectories with no chemotherapy – all trajectories lead to the *S* corner which saturates the tumor. In these figures, solid circles denote the ESS states, while open circles denote the unstable states. Figure 2(b) shows trajectories with *C*_1_ = 0*, C*_2_ = 0.8. In this case, competitive release of the *R*_1_ population allows it to take over the tumor, with all trajectories leading to the *R*_1_ corner. For these parameter values, *R*_1_ is the ESS (and strict Nash equilibrium) of the system. Figure 2(c) with *C*_1_ = 0.8*, C*_2_ = 0 depicts competitive release of the *R*_2_ population, with all trajectories leading to the *R*_2_ corner, showing that *R*_2_ is the ESS of the system.

In figure 2(d) we plot three different constant therapy schedules together, showing how they can intersect at different times. This gives the possibility of switching the therapies off and on at the intersection times in order to create a trajectory that stays in a closed loop and never reaches any of the corners – these closed loops represent scenarios in which the three subpopulations stay in perpetual competition, driven by a time-dependent schedule that shapes the fitness landscapes in such a way as to manage chemotherapeutic resistance. We will describe the systematic construction of these closed loops in §IV. Figure 3 shows the tumor size (using eqn (22)) plotted logarithmically. The untreated tumor (*C*_1_ = 0*, C*_2_ = 0) grows exponentially, while each of the treated tumors initially show tumor regression up until *t* ~ 75, then tumor recurrence due to chemotherapeutic resistance (either of the *R*_1_ or *R*_2_ populations), an unwelcome common scenario. This scenario is clearly depicted in prostate cancer data sets shown in [45] and widely discussed in the clinical literature. Our goal is to show how specially designed adaptive multi-chemotherapies can push the tumor recurrence point further to the right on these plots.

**FIG. 3.**
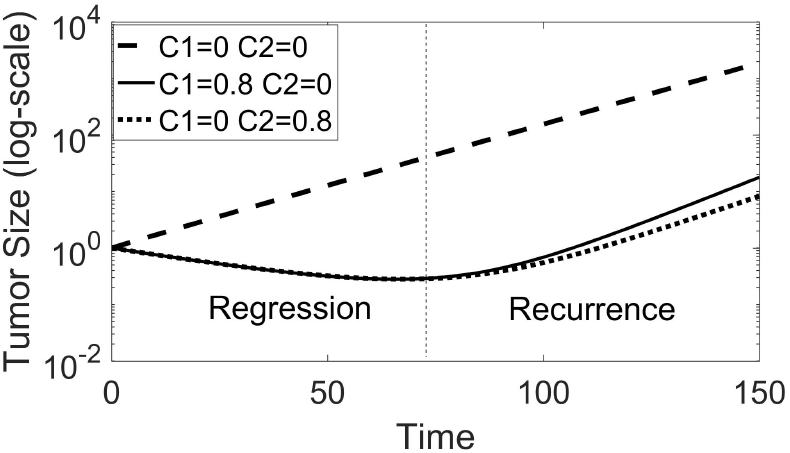
Tumor growth curves (log-plots) for untreated and constant therapies. Tumor recurrence (dashed line) occurs at *t* ∼ 75 dimensionless time units.

The full range of possible profiles are shown in figure 4. For certain ranges of chemo-dosing, there are mixed basins of attraction to each of the corners, hence multiple evolutionary stable states of the system. In the top row, with *C*_1_ = 0, the two critical values where the ESS’s bifurcate are *C*_2_ = 1*/*3 (*R*_1_ changes from unstable to stable via a transcritical bifurcation) and *C*_2_ = 1*/*2 (*S* changes from stable to unstable via a transcritical bifurcation). In the left column, with *C*_2_ = 0, the two critical values where the ESS’s bifurcate are *C*_1_ = 7*/*18 (*R*_2_ changes from unstable to stable via a transcritical bifurcation) and *C*_2_ = 1*/*2 (*S* changes from stable to unstable via a transcritical bifurcation). This panel shows the complexity associated with the stability and basins of attraction of the three tumor subpopulations even in such a simple model, but also gives us the opportunity to exploit the inherent underlying dynamics associated with the different trajectories using piecewise constant chemo-dosing protocols.

**FIG. 4.**
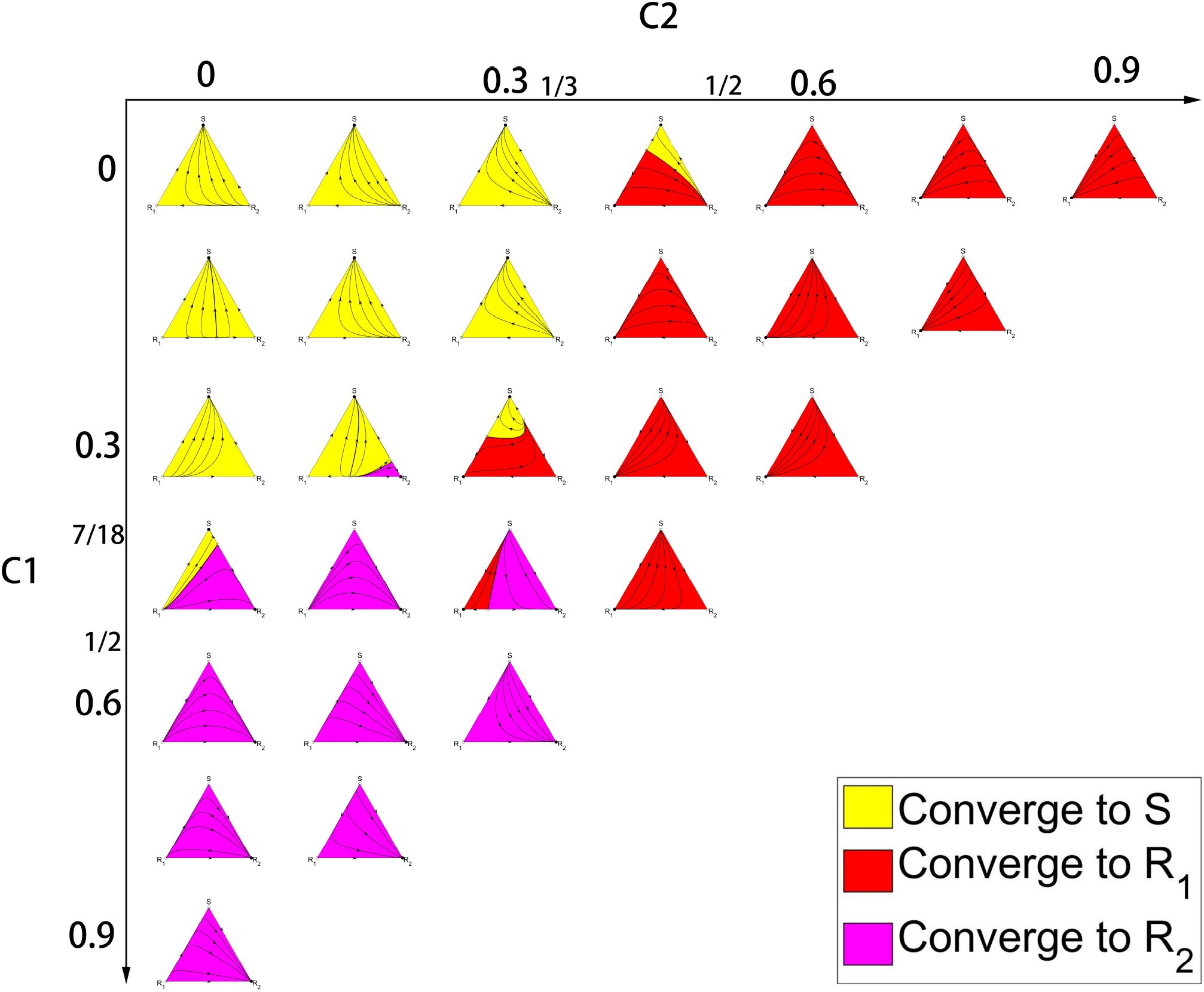
Panel showing the full range of trajectories, ESS and their basins of attraction for constant additive therapies in the range 0 ≤ *C*_1_ ≤ 1, 0 ≤ *C*_2_ ≤ 1, *C*_1_ + *C*_2_ ≤ 1. Bifurcation values along top row are *C*_2_ = 1*/*3, *C*_2_ = 1*/*2. Bifurcation values along left column are *C*_1_ = 7*/*18, *C*_1_ = 1*/*2.

### B. Non-additive interactions

Using the panel in figure 2 as our guide, we focus in more detail on the values *C*_1_ = 0.35, *C*_2_ = 0.27, comparing the effects of synergistic interactions and antagonistic interactions with values *e* = −0.4, −0.3, −0.2, −01., 0, 0.1, 0.2, 0.3, 0.4 in figure 5. For strongly antagonistic interactions (*e* = −0.4, −0.3, −0.2), only the two subpopulations *S* and *R*_2_ compete for dominance (both ESS), with the *R*_1_ population being an unstable state. For strongly synergistic interactions (*e* = 0.2, 0.3, 0.4), the two subpopulations *R*_1_ and *R*_2_ compete for dominance (both ESS), with the population of cells *S* sensitive to both drugs being an unstable state. It is the intermediate regime *e* = −0.1, 0, 0.1 that is the most interesting, and has all three evolutionary stable populations competing for dominance (all three ESS) with intertwined basins of attraction for each. We examine the details of the bifurcations that occur between the two-species co-existence and three-species co-existing states next.

**FIG. 5.**
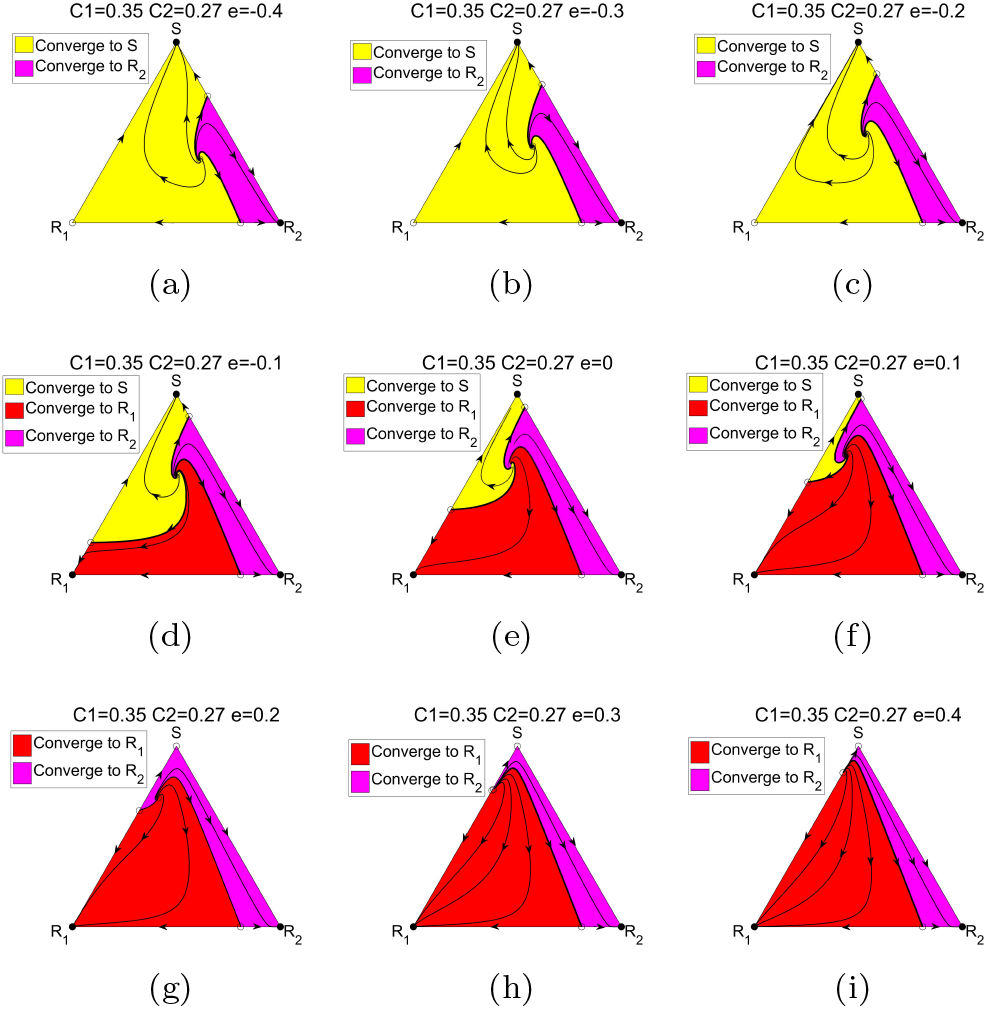
The effect of synergy (*e >* 0) vs. antagonism (*e <* 0) on the ESS and the basins of attraction for a fixed choice of constant combination therapies *C*_1_ = 0.35, *C*_2_ = 0.27. (a) *e* = −0.4; (b) *e* = −0.3; (c) *e* = −0.2; (d) *e* = −0.1; (e) *e* = 0; (f) *e* = 0.1; (g) *e* = 0.2; (h) *e* = 0.3; (i) *e* = 0.4.

#### Transcritical bifurcations

In figure 6, we focus on the range of antagonistic values *e* = −0.21, −0.19, −0.175, −0.15 for fixed values of *C*_1_ and *C*_2_. The relevant bifurcation that occurs at the critical value 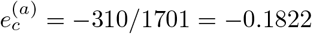 takes place at the *R*_1_ corner, when the fixed point *R*_1_ = 1 goes from un-stable 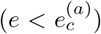 to asymptotically stable 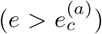 in a transcritical bifurcation (exchange of stability). Through the bifurcation point, a stable fixed point (shown in figure 6(a) outside the triangle below *R*_1_) moves up the *R*_1_ − *S* side of the triangle (*R*_2_ = 0), and exchanges stability with the fixed point at the *R*_1_ corner. In figure 7 we show the details of the collision of eigenvalues that takes place (figure 7(a)) and the process in the *dS/dt* vs. *S* plane (figure 7(b)-(e)). Figure 7(b) shows the classic transcritical bifurcation diagram (see [46]). When the level of antagonism is sufficiently large, there are only the two evolutionary stable states *S* and *R*_2_.

**FIG. 6.**
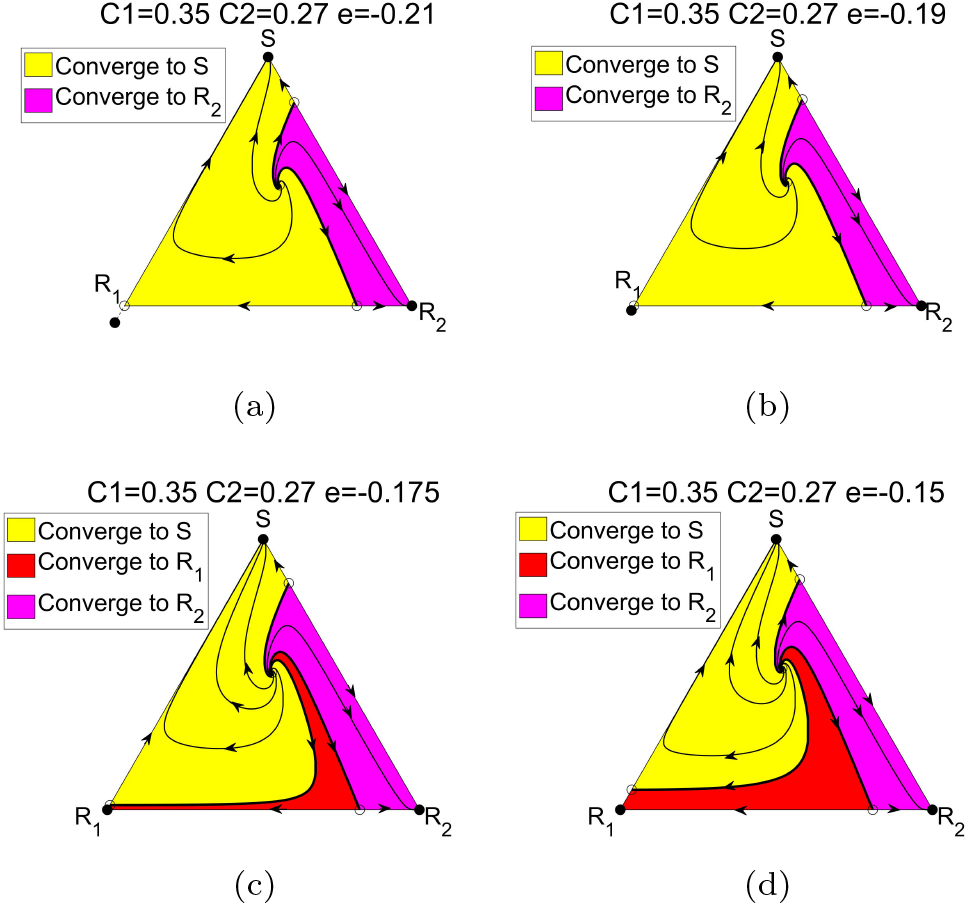
The opening of the basin of attraction for *R*_1_ via a transcritical bifurcation at *e* = −310*/*1701 = −0.1822, *C*_1_ = 0.35, *C*_2_ = 0.27. At the bifurcation point, the areas of the basins of attraction of the respective regions are *S* = 76.2%, *R*_2_ = 13.8%. (a) *e* = −0.21; (b) *e* = −0.19; (c) *e* = −0.175; (d) *e* = −0.15.

**FIG. 7.**
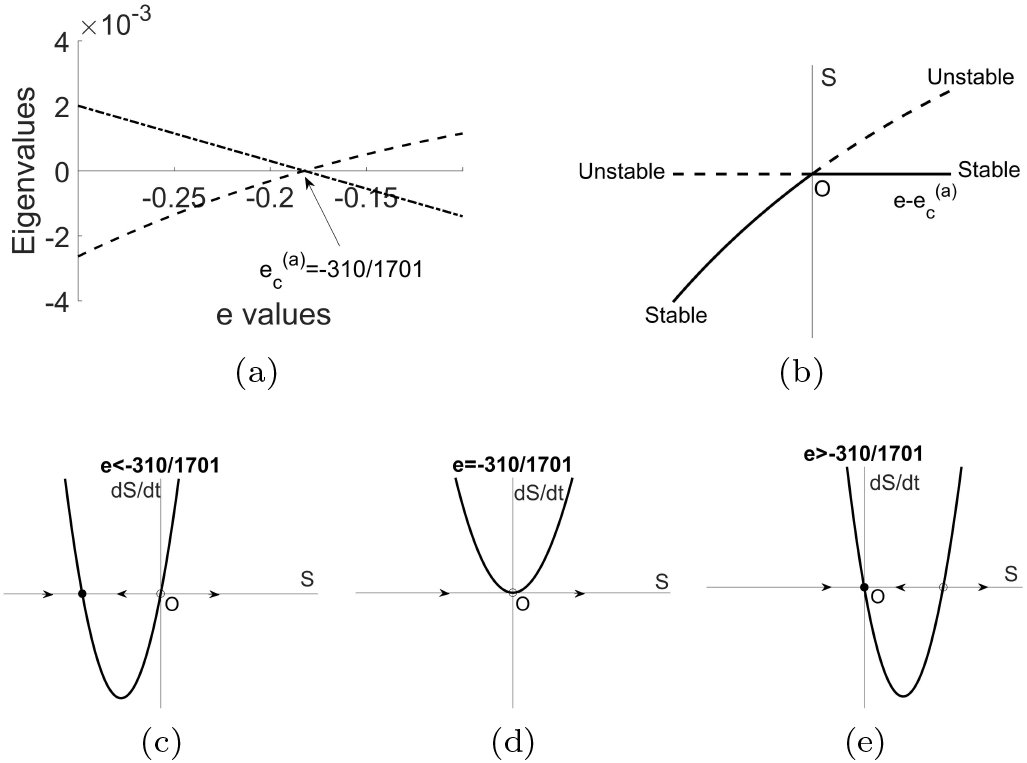
Four ways of depicting the antagonistic transcritical bifurcation. (a) Eigenvalue collision that takes place as the two fixed points collide at the *R*_1_ corner. The other eigen-value remains negative for both fixed points; (b) Transcritical bifurcation diagram; (c) Pre-bifurcation phase plane; (d) Bi-furcation phase plane; (e) Post-bifurcation phase plane.

In figure 8 we highlight the bifurcation that takes place in the synergistic regime around values *e* = 0, 0.1, 0.2, 0.3 for fixed values of *C*_1_ and *C*_2_. The transcritical bifurcation occurs at the critical value 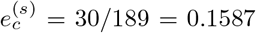 takes place at the *S* corner, when the fixed point *S* = 1 goes from unstable 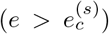 to asymptotically stable 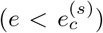. In figure 9 we show the details of the bifurcation diagram that governs this synergistic transcritical bifurcation. When the level of synergism is sufficiently large, there are only the two evolutionary stable states *R*_1_ and *R*_2_.

**FIG. 8.**
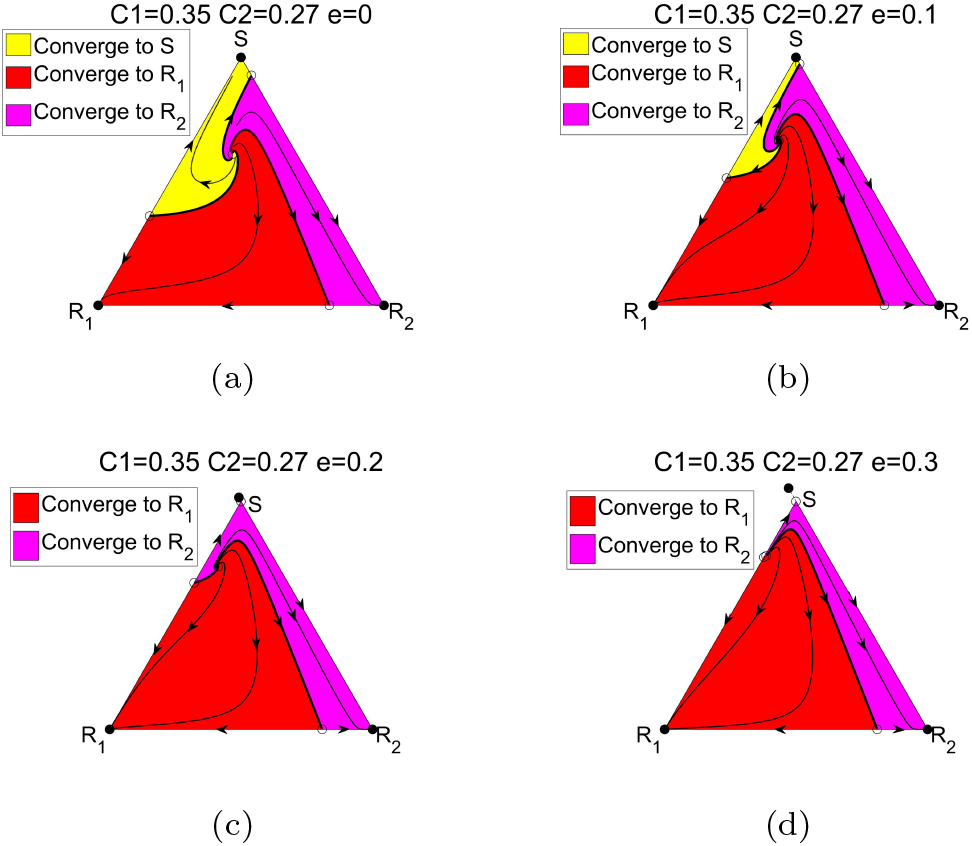
The closing of the basin of attraction for *S* via a transcritical bifurcation at *e* = 30*/*189 = 0.1587, *C*_1_ = 0.35, *C*_2_ = 0.27. At the bifurcation point, the areas of the basins of attraction of the respective regions are *R*_1_ = 72.3%*, R*_2_ = 17.7%. (a) *e* = 0; (b) *e* = 0.1; (c) *e* = 0.2; (d) *e* = 0.3.

**FIG. 9.**
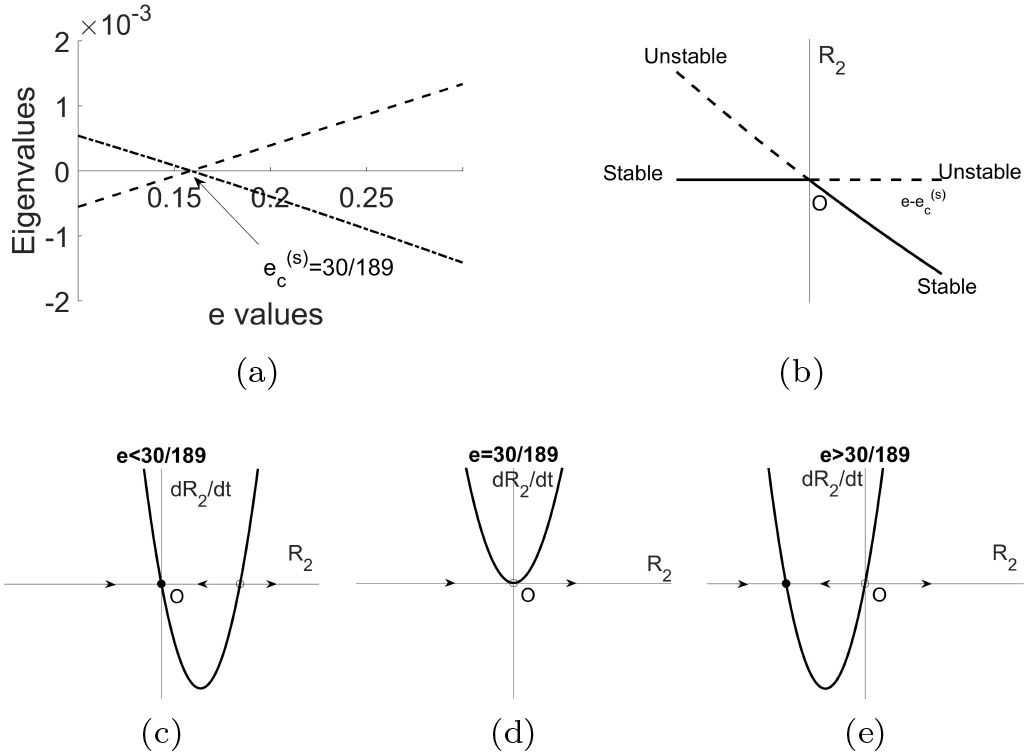
Four ways of depicting the synergistic transcritical bifurcation. (a) Eigenvalue collision that takes place as the two fixed points collide at the *S* corner. The other eigen-value remains negative for both fixed points; (b) Transcritical bifurcation diagram; (c) Pre-bifurcation phase plane; (d) Bi-furcation phase plane; (e) Post-bifurcation phase plane.

In figure 10 we show the areas of three basins of attraction through the full range of values −0.3 ≤ *e* ≤ 0.3. The basin areas begin to rapidly change in the antagonistic regime at *e* = −0.2 and, in general, show much more sensitivity to changes in *e* in the antogonistic regime than the synergistic regime.

**FIG. 10.**
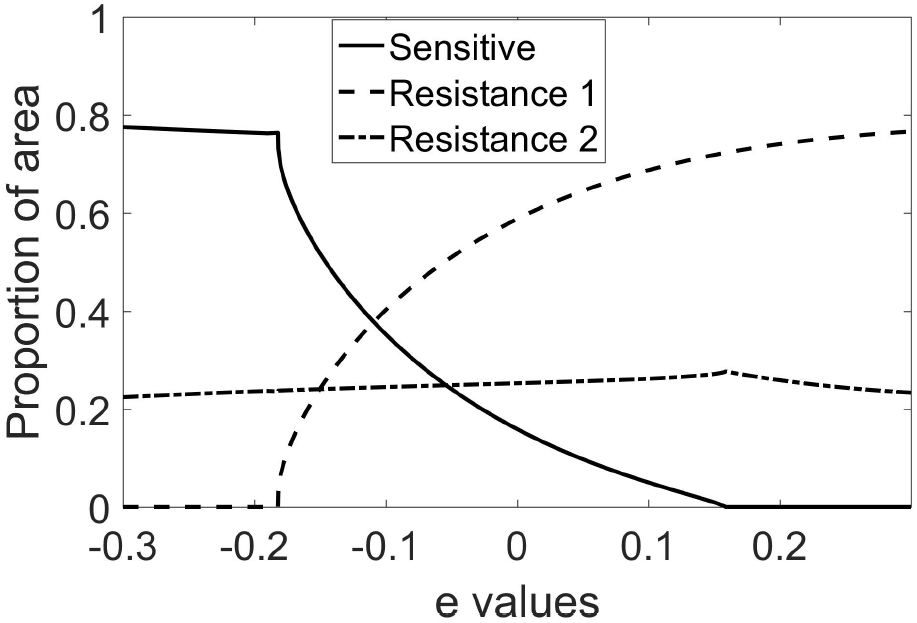
Areas of the three basins of attraction as a function of the parameter *e*. Note the sensitivity of the *S* and *R*_1_ regions in the antagonistic regime −0.2 ≤ *e* ≤ −0.1.

Figure 11 shows the average fitness curves (figure 11(a)) and tumor growth curves (figure 11(b)) through a range of values of *e*. Notice, in the beginning, the average fitness of the antagonistic case *e* = −0.3 is higher, but ends up lower over time. The tumor growth curve in this case follows the same trend - initially tumor growth is highest, then ends up lowest. Tumor recurrence time for the antagonistic case is also pushed later in time (*t* ~ 300) showing the effectiveness of antagonistic drug interactions in terms of managing resistance.

**FIG. 11.**
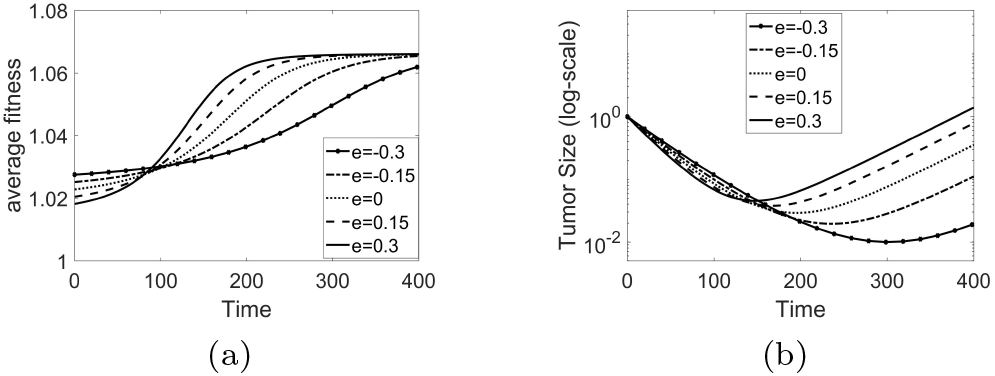
(a) Average fitness curves for *e* = 0.3, 0.15, 0, −0.15, −0.3; (b) Tumor growth curves (log-plot) for *e* = 0.3, 0.15, 0, −0.15, −0.3.

## IV. DESIGNING ADAPTIVE SCHEDULES

Many new possibilities can be created with time-dependent chemoschedules *C*_1_(*t*)*, C*_2_(*t*), if we monitor the balance of the subpopulations and adaptively make changes at judiciously chosen time points. The basic idea is shown in figure 2(d) where we see how the trajectories associated with different constant chemotherapy schedules cross. At any of the crossing times, it is possible to switch from one trajectory to another by switching the values of *C*_1_ or *C*_2_ at those crossing times (typically termed bang-bang control [47]). This basic procedure allows us to design schedules that take us from any point *A* in the triangle to any other point *B* along the legs of a path that are separated by the switching times. Using multiple time-switching, we can also design trajectories that form closed loops and never converge to any of the corners.

Figure 12 shows examples of how closed (piecewise differentiable) orbits are designed in practice. In each of the figures, point O is fixed, as is the untreated (blue) curve with *C*_1_ = 0, *C*_2_ = 0. Consider the loop OABO created figure 12(a), with additive interaction parameter *e* = 0. In traversing the OA leg, we use *C*_1_ = 0.5, *C*_2_ = 0.2. When the trajectory reaches point A, we switch to *C*_1_ = 0, *C*_2_ = 0, i.e. no therapy. When we reach point B, we switch to *C*_1_ = 0.2, *C*_2_ = 0.5 until we reach point O, and then we start the schedule again to traverse the same loop as many times as we desire. The dose density plot is shown in figure 12(b). Figures 12(c),(d) uses the same dosing values, but with *e* = 0.3 (synergistic). The loop in this case is larger (encloses more area), so there is a larger deviation in the sub-populations throughout the loop than there was for the additive case. Figure 12(e),(f) shows an example of an antagonistic *e* = −0.3 adaptive therapy loop. Table I shows the total dose *D*, time period *T*, and average dose *D/T* for each case. The antagonistic adaptive loop delivers the smallest total dose over the shortest time-period to achieve one closed loop, whereas the synergistic adaptive loop delivers the smallest average dose over the loop, since the time-period is longest.

**FIG. 12.**
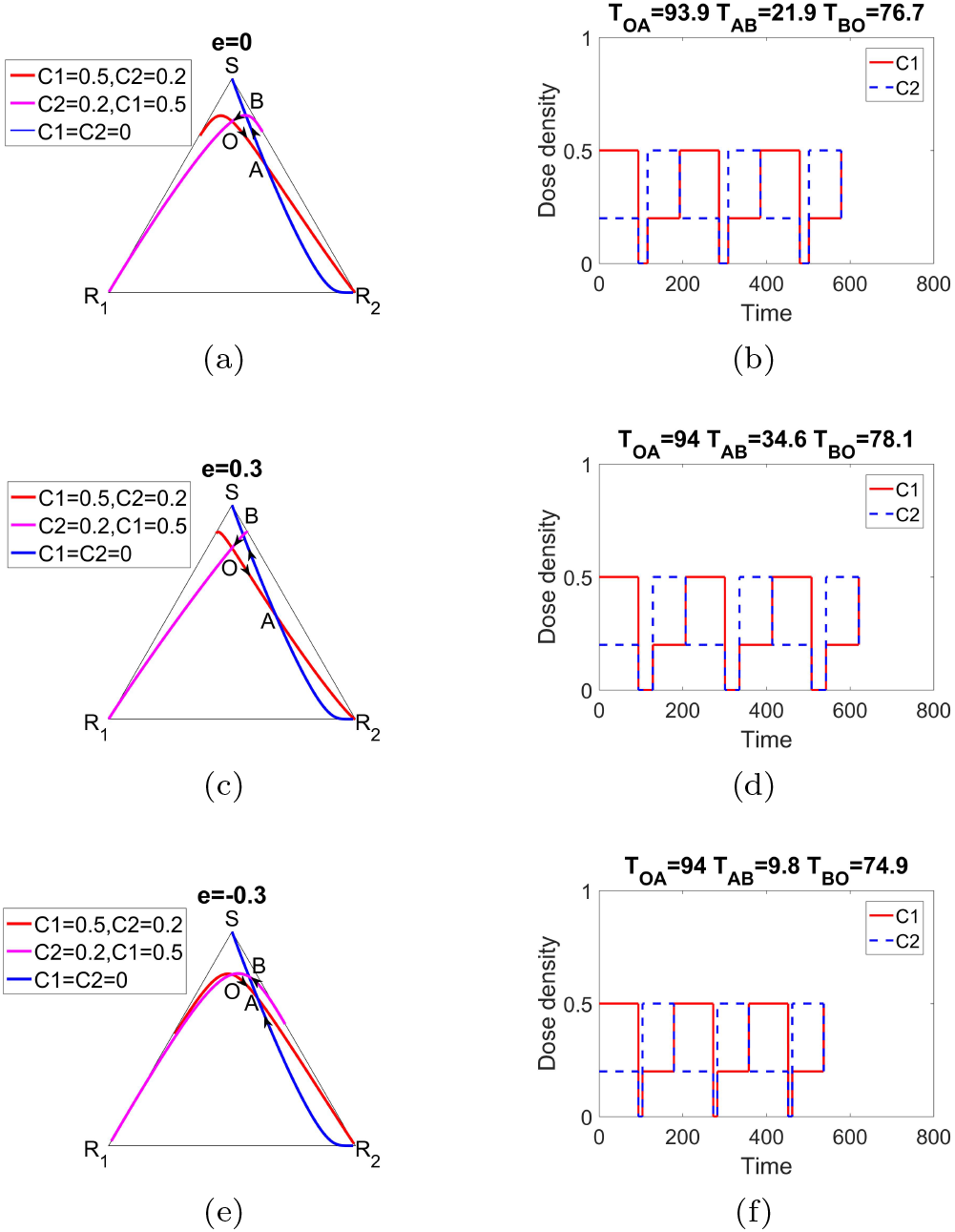
Closed loop adaptive schedules *OABO* using three cycles for *e* = 0, 0.3, −0.3. (a)*e* = 0, *OA*: *C*_1_ = 0.5*, C*_2_ = 0.2, *AB*: *C*_1_ = 0*, C*_2_ = 0, *BO*: *C*_1_ = 0.2*, C*_2_ = 0.5; (b) Corresponding adaptive schedule; (c) *e* = −0.3, *OA*: *C*_1_ = 0.5*, C*_2_ = 0.2, *AB*: *C*_1_ = 0*, C*_2_ = 0, *BO*: *C*_1_ = 0.2*, C*_2_ = 0.5; Corresponding adaptive schedule; (e) *e* = −0.3, *OA*: *C*_1_ = 0.5*, C*_2_ = 0.2, *AB*: *C*_1_ = 0*, C*_2_ = 0, *BO*: *C*_1_ = 0.2*, C*_2_ = 0.5; (f) Corresponding adaptive schedule.

**TABLE I.** Total dose (D), Time period (T), and Average Dose (D/T) associated with adaptive therapies with antagonistic, additive, and synergistic drug interactions.

| $e$ | -0.3 | 0 | 0.3 |
| --- | --- | --- | --- |
| **D** | 118.23 | 119.42 | 120.47 |
| **T** | 178.7 | 192.5 | 206.7 |
| **D/T** | 0.6616 | 0.6204 | 0.5828 |

Figure 13 shows the tumor growth curves for each of the adaptive schedules as compared with the untreated growth curve (exponential growth), and constant therapy curve (each shows tumor regression followed by recurrence). In each case, the adaptive loop overcomes recurrence, with the antagonistic schedule minimizing the tumor re-growth leg (AB) of the schedule.

**FIG. 13.**
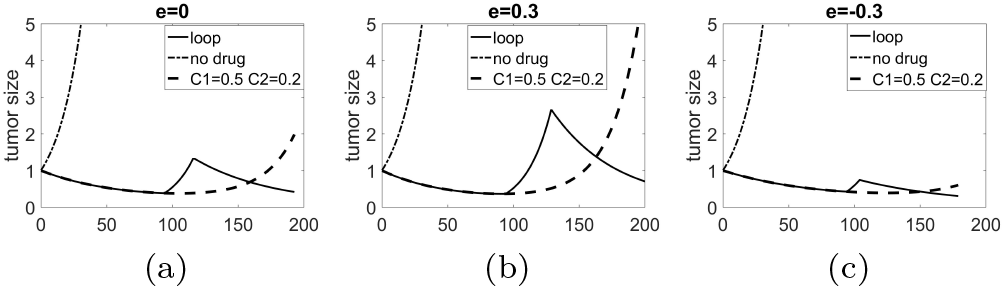
Tumor growth curves for the adaptive schedules from figure 12 as compared with untreated growth (exponential) and constant schedule which eventually leads to tumor recurrence. (a) e = 0; (b) e = 0:3; (c) e = −0:3.

A final metric that we use for comparisons is the rate of adaptation of the subpopulation *i*, defined by:

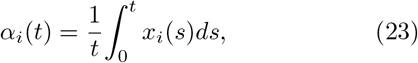

which is the time-average of subpopulation level *x_i_*(*t*).

Figure 14 shows the rates of adaptation associated with the two resistant subpopulations *R*_1_ and *R*_2_ during the course of the adaptive schedules. Notice the rate of adaptation is lowest for the antagonistic interaction, which is the main reason the tumor growth curve in figure 13(c) is most effective at controlling and delaying tumor recurrence as the cell population evolves.

**FIG. 14.**
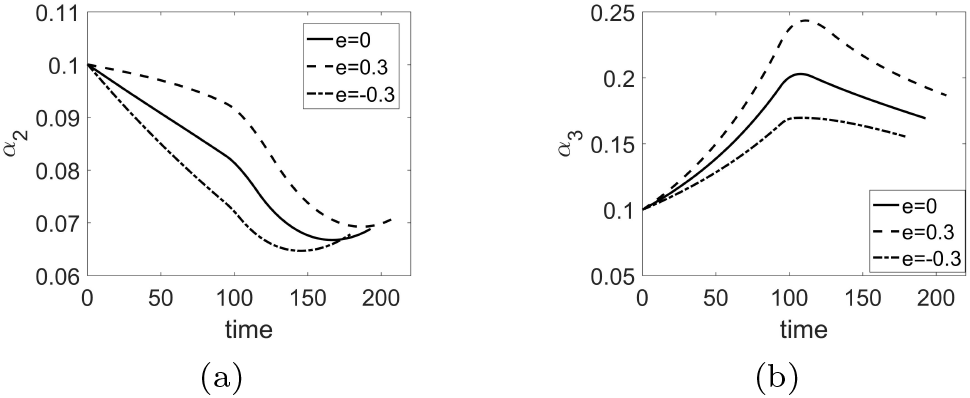
Rates of adaptation for *e* = 0, 0.3, −0.3: (a) *R*_1_ rate of adaptation; (b) *R*_2_ rate of adaptation; (c) *x*_1_ rate of adaptation; (d) *x*_2_ rate of adaptation;

## V. DISCUSSION

The role of synergistic vs. antagonistic combination drug interactions on the dynamical balance of evolving subpopulations of cells is not simple to characterize. In general terms, antagonistic interactions are able to exert a more targeted and subtle range of influences on an evolving population than synergistic ones which, roughly speaking, cause the two drug combination to effectively act as one. This limits their flexibility for designing effective strategies to manage resistance, although might be useful in producing a bigger killing effect with a smaller dose. As a by-product of the ability to deliver a more nuanced effect on the balance of cells, the sensitivity to small changes in the relative doses of the two drugs in an antagonistic setting seems to be higher than in a synergistic setting (as shown best in figure 10). This, perhaps, makes these interactions harder to control. Tumor recurrence times for the antagonistic drug interactions are delayed more effectively than for synergistic interactions, consistent with the fact that adaptation rates are slower for antagonistic interactions. Anatagonistic drug inter-actions provide more flexibiility in designing adaptive re-sistance management schedules. Additional features to our model could be the introduction of mutations that might occur in response to the chemotherapy, which potentially could be handled using a mutational replicator system [3], or a finite-cell Moran process based model [40]. It might also be possible to analyze existing individual patient data and tumor response curves (as was done in [45]) to design and optimize better multi-drug strategies retrospectively, which is ongoing by our group. There are also ongoing adaptive multi-drug clinical trials at the Moffitt Cancer Center that show promise in prostate cancer patients.

## ACKNOWLEDGMENTS

We gratefully acknowledge support from the Breast Cancer Research Foundation (BCRF), the Jayne Koskinas & Ted Giovanis Foundation (JKTG), as well as the Army Research Office MURI Award #W911NF1910269 (2019-2024).

## References

[1] P. Newton and Y. Ma, Phys. Rev. E 99 (2019).

[2] J. Hofbauer and K. Sigmund, Evolutionary Games and Population Dynamics (Cambridge University Press, 1998).

[3] M. A. Nowak, Evolutionary Dynamics (Harvard University Press, 2006).

[4] R. Axelrod, D. E. Axelrod, and K. J. Pienta, Proc. Natl. Acad. Sci. 103, 13474 (2006).

[5] J. West, Z. Hasnain, J. Mason, and P. Newton, Converg. Sci. Phys. Oncol. 2, 035002 (2016).

[6] R. Gatenby, A. Silva, R. Gillies, and B. Frieden, Cancer Research 69, 4894 (2009).

[7] R. Gatenby and J. Brown, Cold Spring Harbor Perspectives in Medicine, doi: 10.1101/cshperspect.a033415 (2017).

[8] A. Silva and R. Gatenby, Biology Direct 5, 1 (2010).

[9] P. Enriquez-Navas, J. Wojtkowiak, and R. Gatenby, Cancer research 75, 4675 (2015).

[10] J. A. Gallaher, P. M. Enriquez-Navas, K. A. Luddy, R. A. Gatenby, and A. R. Anderson, (2017).

[11] J. Zhang, J. Cunningham, J. Brown, and R. Gatenby, Nature Comm. 8, 1816 (2017).

[12] J. Foo and M. F, J. Theor. Bio. 263, 179 (2010).

[13] J. Foo and M. F, J. Theor. Bio. 355, 10 (2014).

[14] R. Gillies, D. Verduzco, and R. Gatenby, Nature Reviews Cancer 12, 487 (2012).

[15] I. Bozic and M. Nowak, Ann. Rev. of Can. Bio. 1, 203 (2017).

[16] O. Lavi, M. Gottesman, and L. D, Drug Res. Updates 15, 90 (2012).

[17] N. Komarova and D. Wodarz, Proc. Natl. Acad. Sci. 102, 9714 (2005).

[18] M. Hegreness, N. Shoresh, D. Damiann, D. Hartl, and R. Kishony, Proc. Nat’l Acad. Sci. 105, 13977 (2008).

[19] C. Bliss, Annals of Applied Biology 26, 585 (1939).

[20] S. Loewe, Arzneim. Forsch. 3, 285 (1953).

[21] T. Chou and D. Rideout, Synergism and Antagonism in Chemotherapy (Academic Press, NY, 1991).

[22] P. Sudalagunta, M. Silva, R. Canevarolo, R. Gatenby, G. R, R. Baz, M. Meads, K. Shain, and A. Silva, EBioMedicine 54, 102716 (2020).

[23] A. Read, T. Day, and S. Huijben, Proc. Natl. Acad. Sci. 108, 10871 (2011).

[24] D. Basanta, R. Gatenby, and A. Anderson, Molecular pharmaceutics 9, 914 (2012).

[25] J. West and P. Newton, Cancer Research, doi: 10.1158/0008 (2017).

[26] J. West, Y. Ma, and P. Newton, Cancer Research 80, 1578 (2020).

[27] D. Basanta and A. Anderson, Interface Focus 3, 20130020 (2013).

[28] J. Michel, P. Yeh, R. Chait, R. Moellering, and R. Kishony, Proc. Nat’l Acad. Sci. 105, 14918 (2008).

[29] D. Andersson, N. Balaban, F. Baquero, P. Courvalin, P. Glaser, U. Gophna, R. Kishony, S. Molin, and T. Tonjum, FEMS Microbiology Rev. (2020).

[30] D. Russ and R. Kishony, Nature Microbiology 3, 1339 (2018).

[31] M. Baym, L. Stone, and R. Kishony, Science 351, DOI: 10.1126/science.aad3292 (2016).

[32] S. Kim, T. Lieberman, and R. Kishony, Proc. Nat’l Acad. Sci. 40, 14494 (2014).

[33] K. Wood, K. Wood, S. Nishida, and P. Cluzel, Cell Reports 6, 1073 (2014).

[34] D. Nichol, P. Jeavons, A. Fletcher, R. Bonomo, P. Maini, J. Paul, R. Gatenby, A. Anderson, and J. Scott, PLoS Comp. Bio. 11(2015).

[35] N. Yoon, R. Vander Velde, A. Marusyk, and J. Scott, Bull. Math. Bio. 80, 1776 (2018).

[36] M. Lipsitch and B. Levin, Antimicrobial Agents and Chemotherapy 41, 363 (1997).

[37] I. Bozic and M. Nowak, Proc. Natl. Acad. Sci. 111, 15964 (2014).

[38] S. Hummert, K. Bohl, D. Basanta, A. Deutsch, S. Werner, G. Theiben, A. Schroeter, and S. Schuster, Molecular BioSystems 10, 3044 (2014).

[39] K. Bohl, S. Hummert, S. Werner, D. Basanta, A. Deutsch, S. Schuster, G. Theiben, and A. Schroeter, Molecular BioSystems 10, 3044 (2014).

[40] J. West, Z. Hasnain, P. Macklin, and P. Newton, SIAM Review 58, 716 (2016).

[41] A. Traulsen, J. C. Claussen, and C. Hauert, Physical Review Letters 95, 238701 (2005).

[42] B. Wu, P. Altrock, L. Wang, and A. Traulsen, Phys. Rev. E 82, 046106 (2010).

[43] B. Walsh and M. Blows, Annu. Rev. Ecol. Evol. Syst. 40, 41 (2009).

[44] J. West and P. Newton, Proc. Nat’l Acad. Sci. 116, 1918 (2019).

[45] J. West, Y. Ma, and P. Newton, J. Theor. Bio. 455, 249 (2018).

[46] S. Strogatz, Nonlinear Dynamics and Chaos (Westview Press, 2015, 2nd Ed.).

[47] U. Ledzewicz and H. Schattler, J. Optimization Theory and Appl. 114, 609 (2002).

